# A newly developed whole genome sequencing protocol enables early tracking of Enterovirus D68 molecular evolution

**DOI:** 10.1101/2024.09.30.615419

**Authors:** Federica AM Giardina, Greta Romano, Guglielmo Ferrari, Laura Pellegrinelli, Arlinda Seiti, Cristina Galli, Elena Pariani, Antonio Piralla, Fausto Baldanti

## Abstract

Background Human enterovirus D68 (EV-D68) has been associated with an increase in mild-to-severe pediatric respiratory diseases worldwide. The rate of circulation of this virus is largely underestimated in the population and genetic evolutionary data are usually available only for partial sequences (e.g VP1 region). To achieve a timely genomic surveillance, a reliable, high-throughput EV-68 sequencing assay is required. Here we report an improved high-throughput EV-D68 whole-genome sequencing (WGS) assay performed directly on clinical samples that is suitable for short-read sequencing platforms. Between June and December 2022, a total 37 (1.9%) respiratory samples were EV-D68 positive and together with 52 additional samples with a median cycle threshold of 28.3, ranging from 18 to 36.8 were included in the validation analyses. Overall, all the primers had good performance and no mismatches were detected in more than 85% of sequences (932 WGS dataset). Using a cut-off of Cq <32 in at least 85.5% of samples a WG or PG was obtained, confirming an acceptable positive sequencing rate for the designed method. A total of 65 WGS were obtained and have a mean coverage of 98.4% across the genome, with a median depth of 6158x (range 2815x-7560x). Based on the obtained data, this method is cost effective resulting in an easy-to-perform protocol helpful for tracing the evolution of EV-D68 in protein different from VP1. EV-D68 could become a significant pathogen for public health in the next future, and thus this protocol for whole genome sequencing could help clinical and molecular virologists to be ready for molecular epidemiology surveillance.

## 1. Introduction

Enterovirus D68 (EV-D68) belongs to the *Picornaviridae* family, genus Enterovirus. It was isolated in 1962 for the first time from four children with severe respiratory syndromes (Schieble *et al*., 1967). Since its first detection, EV-D68 cases were very sporadic until 2014, when it caused a very large outbreak in the United States and Canada first (CDC, 2014; Midgley *et al*., 2014) and all over Europe later with thousands of cases detected (Bragstad *et al*., 2015; Esposito *et al*., 2015; Gimferrer *et al*., 2014). EVD68 is associated with respiratory illness, ranging from mild to severe, especially in children with underlying respiratory conditions such as asthma or pneumonia (Schuster *et al*., 2015). Occasionally, EV-D68 infection could have neurological complications with symptoms of acute flaccid myelitis or acute flaccid paralysis (AFM/AFP) (Suresh *et al*., 2018). So far, the emergence of new EV-D68 subclades has been traced by sequencing of VP1 gene, which is one of the main proteins composing viral capsid (Piralla *et al*., 2018). Furthermore, in the last years, a huge effort on whole-genome sequencing (WGS) of EV-D68 has been showed by several studies (Bhardwaj et al., 2019; Stelzer-Braid et al., 2022). The co-circulation of several EV-D68 subclades has evidenced the needed of monitor ED-D68 evolution not only in VP1 region (Piralla et al., 2023). In addition, WGS allow to identify molecular patterns potentially associated with uncommon clinical syndromes (Graphin et al.,2023; Piralla et al., 2023; Singanayagam et al., 2023). Finally, the study of WGS provide new insight the emergence of recombinant strains also previously observed for enterovirus species A-C (Lukashev et al., 2014; Muslin et al., 2019; Smura et al., 2014). The present study aimed to: i) design a new PCR-based next-generation sequencing (NGS) assay for WGS of EV-D68, ii) assess the quality of primers using a phylo-primer-missmacth analysis and iii) sequence EV-D68 strains circulated in the Northern Italy during the 2022-2023 winter season and, iv) evaluate the viral evolution.

## 2. Material and Methods

### 2.1. Primer design ad in-silico evaluation

A total of 932 EV-D68 complete genome sequences, with more than 7200 nucleotides (nt) were retrieved from online repositories (GenBank accessed on 16 September 2023) and aligned with MAFFT v7.525 (Katoh and Standley, 2013). Primers for RT-PCR were designed based on conserved regions of EV-D68 in silico PCR using FastPCR software (Kalendar et al., 2017). The specificity of these primers was further checked by BLAST to ensure that the primers were specific for EV-D68. Three pairs of primers were designed to amplify 3 amplicons. Degenerate bases were introduced in the primers to cover the great majority of variants. The primers efficiencies and the theoretical annealing temperature were evaluated using the FastPCR software (Kalendar et al., 2017). A list of selected primer is reported in Table 1. Similar to the approach used for primer design, representative EV-D68 sequences were obtained from GenBank and primers sequences were mapped against the alignment to analyse the number of mismatches per strain (phylo-primer-missmacth analysis). A Maximum likelihood (ML) tree was constructed in IQ-TREE5 (Minh et al., 2020) with a substitution model chosen according to BIC within the IQ-TREE internal pipeline with 1000 bootstrap replicates. The mismatches were tabulated for visualization and overlayed with the phylogenetic tree as previously reported (Fan NG et al. 2023) using in-house script developed by R programming language (https://www.R-project.org/).

**Table 1.**
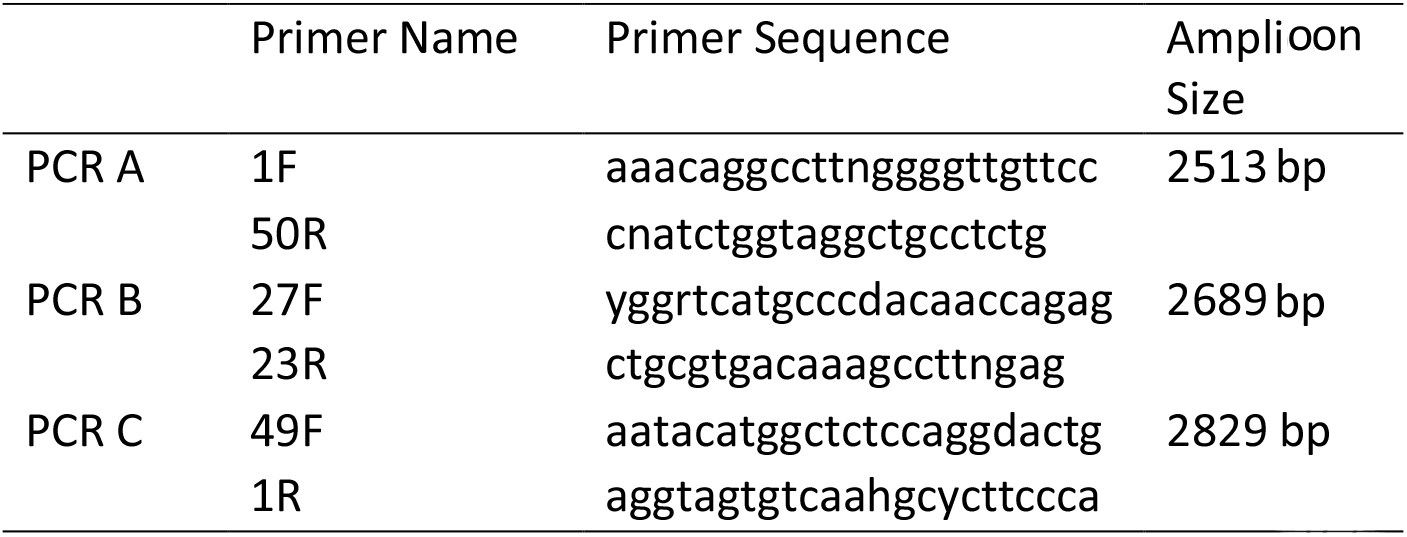
Primer pairs and amplicon size for EV-D68 whole genome amplification.

### 2.2. Molecular screening for EV-D68

All respiratory samples (nasal swabs and bronchoalveolar lavages) collected from June to December 2022 at Microbiology and Virology Department, Fondazione IRCCS Policlinico San Matteo, Pavia, Italy, were included in this study. Nucleic acid extraction was performed using EZ1&2™ Virus Mini Kit v2.0 on platform EZ1 Advanced XL (Qiagen, Heidelberg, Germany) and tested for a panel of respiratory viruses as previously reported (Piralla *et al*., 2011). Residual RNA samples were tested for the presence of EV-D68 using pools of five samples using an EV-D68 specific real-time according to published protocols (Piralla *et al*., 2015). Results were given as cycle of quantification (Cq) values that inversely reflects viral load. A Cq ≥ 40 was set as negative cut-off. Briefly, the assay was performed using QuantiFast®Pathogen RT-PCR+IC Kit (Qiagen, Heidelberg, Germany) and carried out on Rotor-Gene Q thermal cycler, with the following thermal profile: 50°C for 20 minutes, 95°C for 5 minutes, 45 cycles at 95°C for 15 seconds and 60°C for 30 seconds. EV-D68 positive pools were then split and single samples were tested with the same protocol. In addition, to validate the EV-D68-WGS protocol, a total of 52 additional EV-D68 positive samples identified at the Biomedical Sciences for Health Department, University of Milan in the period 2018-2022 were included in the analysis (Pellegrinelli et al., 2019).

### 2.3. Whole genome amplification and sequencing

SuperScript IV One-Step RT-PCR System with ezDNase (Invitrogen) was used for EV-D68 whole genome amplification with three different primer sets designed for this study (Figure 1 and Table 1). Three segments of 2513 bp, 2689 bp and 2829 bp, respectively with overlapping region of 334 and 344 nt were designed. Amplicons were processed using the same thermal profile: 55°C for 10 minutes, 98°C for 2 minutes; 45 cycles at 98°C for 10 seconds, 60°C for 10 seconds, 72°C for 2 minutes with final elongation was at 72°C for 5 minutes. The amplicon sizes were evaluated by gel electrophoresis. Amplicons were purified with AMPure XP beads and eluted in 50 µl 1X TE buffer and quantified using Qubit dsDNA BR Assay Kit and QubitTM 4 Fluorometer (ThermoFisher). Equal amounts of RT-PCR products from three amplicons were pooled together for downstream NGS library preparation and sequencing. Genomic libraries were prepared using Nextera XT Library Preparation Kit (Illumina, San Diego, CA, USA), according to the manufacturer’s instructions. Libraries were quantified with Qubit 1X dsDNA HS Assay kit, normalized at the same concentration, and then pooled together. The pool was denatured with 0.2 N NaoH and then diluted to a final concentration of 10 pM. Sequencing was performed on MiSeqDx platform using MiSeq Reagent kit V2 (Illumina, San Diego, CA, USA). Sequences obtained were analyzed on INSaFLU (Borges *et al*., 2018), a user-friendly bioinformatic web-based tool that deals with primary sequencing data. EV-D68 genome with accession number OP267512.1 was used as the reference strain. e

**Figure 1.**
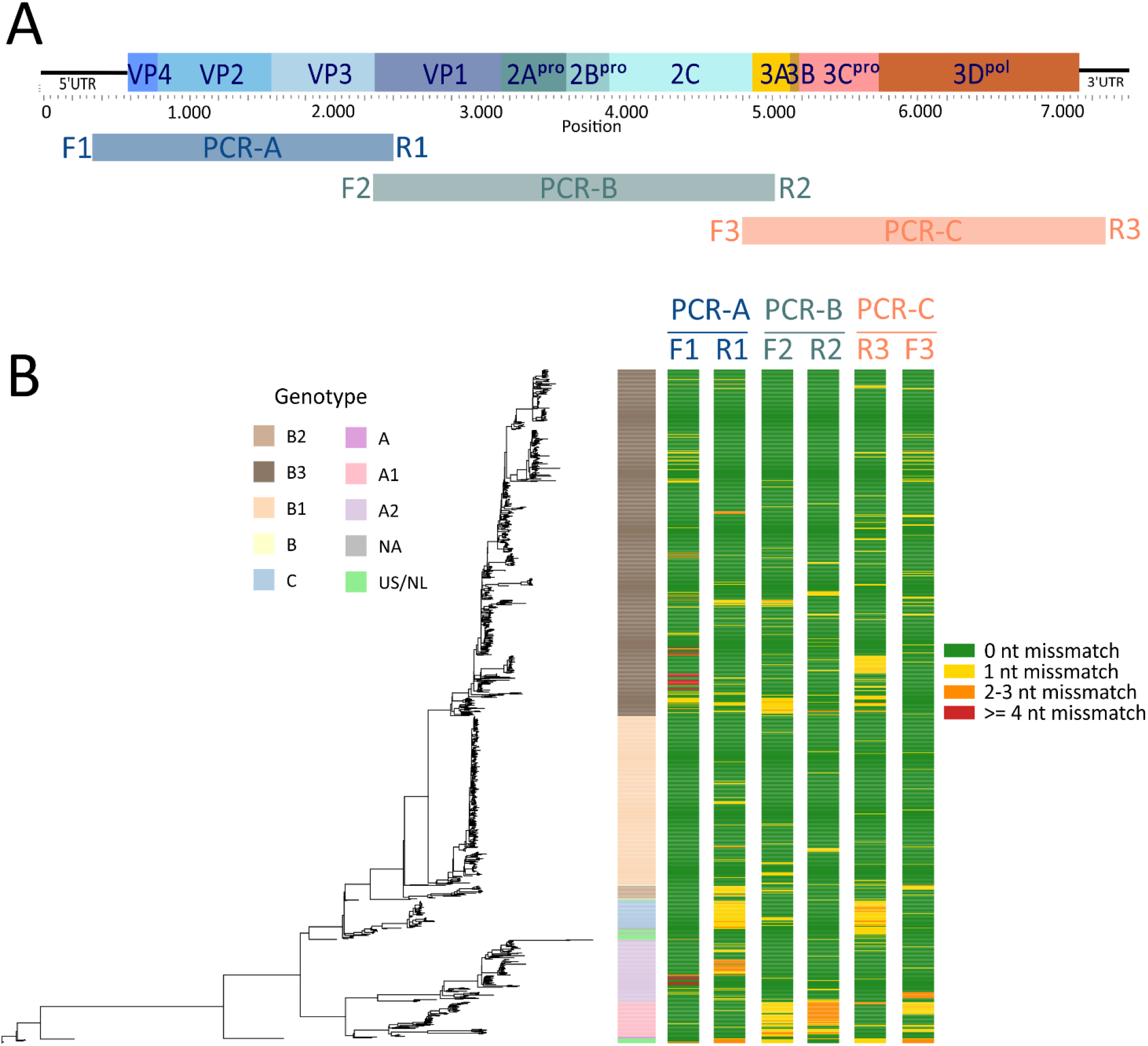
**A)** PCR design for EV-D68 complete genome sequencing. A) Viral genome was divided into three segments, overlapping by 334 and 344 nt respectively. For each segment, primers used and length of amplicon produced are listed. B) Phlyo-primer-mismatch graph for visualization of primer/probe mismatches against th EV-D68 phylogeny. Alignment of 359 representative deduplicated VP1 nucleotides were use to construct a neighbor-joining tree. Primer/probe mismatches to each sequence were tabulated and overlayed with the tree. Sense primer, antisense primer, and probe were evaluated in this order for each assay. WU and NU assays have two antisense primers. S, sense primer; A, antisense primer; A1, antisense-1 primer; A2, antisense-2 primer; P, probe Blue, 0 mismatch; Yellow, 1 mismatch; Orange, 2 mismatches; Red, 3 or more mismatches.

### 2.4. Phylogenetic analyses

The alignment and phylogenetic analysis were performed with aligned with MAFFT v7.525 (Katoh and Standley, 2013) and IQ-TREE v.2.2.2.6 (Minh et al., 2020), respectively. Figures showed in this article were created using Rstudio Computer Software v0.98.1074 (RStudio Team, 2015 RStudio: Integrated Development for R. RStudio, Inc., Boston, MA; http://www.rstudio.com). We integrated all the EV-D68 Italian genomes originated in this study with 963 sequences available from GenBank including metadata (e.g., sample collection date, country). Enterovirus Genotyping Tool (Kroneman et al., 2011) was used to derive the enterovirus genotype of the collected dataset.

### 2.5. Statistical Analysis

Comparisons between two groups were performed using the two-tailed t-test. P values less than 0.05 were considered as statistically significant. R software and packages were used for making graphs and curve fitting analyses. All the statistical analyses were performed using GraphPad Prism software version 8.3 (GraphPAD Software, San Diego, CA).

## 3. Results

### 3.1. PCR primer design in silico

To validate the PCR primers used in this study we performed an in-silico analysis to evaluate the performances capture for all the genotypes. Six PCR primers were designed in the more conserved region of all EV-D68 known genotypes overlapping all the genes from 5’UTR to 3’UTR, as described below (see methods section 2.1). Figure 1 showed the number of mismatches per primer in correspondence of the position inside the tree for all EVD-68 sequences included in the dataset. Overall, all the primers had good performance mainly highlighted by green color where no mismatches were detected. In detail, among the forward primers, in 940 out of 1027 (91.5%) sequences no mismatches were detected in 1F primer, followed by 49F (927/1027; 90.3%) and 27F (901/1027;87.7%). However, in 30 EV-D68 sequences belonged to genotype A2 and B3 more than four mismatches in primer 1F were observed. Regarding reverse primers, 23R had the best performance with 936 out of 1027 (91.1%) EV-D68 sequences with no mismatches followed by 1R (877/1027; 85.4%) and 50R (876/1027; 85.3%), respectively.

### 3.2. EV-D68 Positive samples and WGS results

A total of 1932 respiratory samples were collected between June and December 2022 and were screened for the presence of EV-D68 genome. Among these, 37 (1.9%) were EV-D68 positive. Overall, to validate the new amplicon-based whole genome sequencing assay, complete genome amplification was performed on all 37 EV-D68 positive samples and in 52 additional samples for a total of 89 EVD-68-positive samples with a median cycle threshold of 28.3, ranging from 18 to 36.8. The three PCRs were successful for 83/89 (93.2%) samples while in 6/89 (6.8%) no amplicons were obtained. In addition, for 11/89 (12.4%) samples PCR-positive no good quality sequencing data were obtained and for 6/89 (6.7%) samples only partial genome (mean 5668 nt; range 4935-6593 nt) was obtained. Overall, a complete genome sequence was obtained in 73.0% of samples (65/89) with a median length of 7349 nt (range 7347 to 7356 nt). A complete overview of sequencing data for each sample are reported in supplementary data (Table S1). As showed in Figure 2A, samples in which WG was obtained had viral load higher than samples without sequencing results (mean 26.7 *vs* 28.9 Cq; p=0.04). Overall, using a cut-off of Cq <32 in at least 85.5% of samples a WG or PG was obtained, confirming an acceptable positive sequencing rate for the designed method.

**Figure 2.**
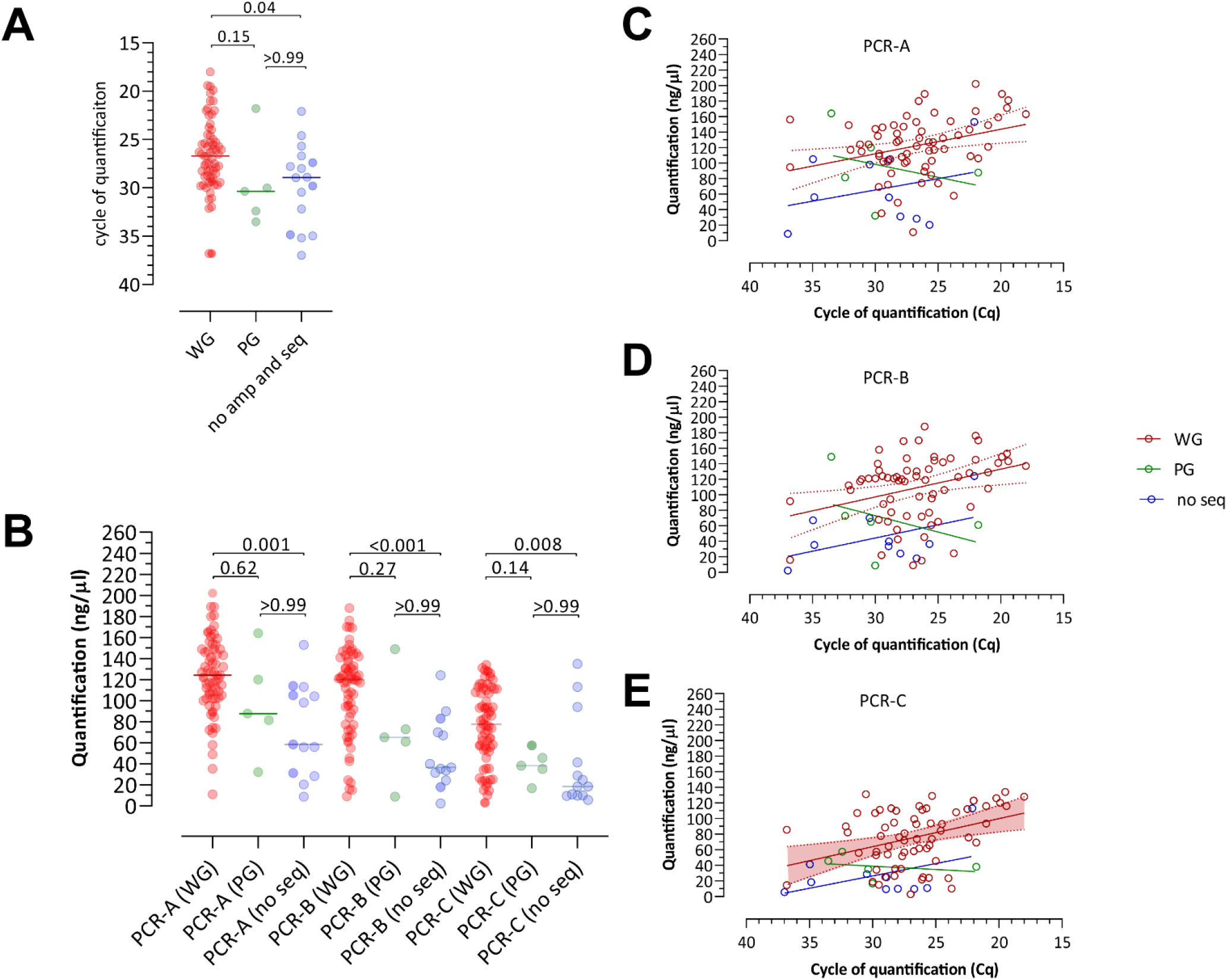
Comparison of EV-D68 WGS protocol performances.

Among 83/89 (93.3%) samples with positive amplicon results the quantification of EV-D68 cDNA was higher for all the three amplicons in samples with WG as compared to samples without sequencing results (Figure 2B; p<0.05). On the same time, there was a correlation between the viral load and the quantification of amplicons obtained for all the three amplicons in WG and samples without sequencing results (Figure 2C-E).

A total of 65 WGS were obtained and have a mean coverage of 98.4% across the genome, with a median depth of 6158x (range 2815x-7560x). The median percentage of genome covered by at least 1-fold was 98.4% (range 98.1%-98.5%) while the median percentage of genome covered by at least 10-fold was 98.3% (range 77.6%-98.4%). In 5/89 EV-D68 positive samples (5.6%) a partial genome sequence was obtained. For these samples, the median length of genome obtained was 5710 nt (range 4935-6593 nt). The mean depth of coverage was 2577x (range 179x-5310x; Figure S1). The median percentage of genome sequence covered by at least 1-fold was 95.6% while the median percentage of genome covered by at least 10-fold was 75.2% (range 90%-97.7% and 57.8%-91.6%, respectively).

### 3.3. Phylogenetic analysis

Figure 3 shows the phylogenesis of the dataset which revealed 3 major clades A, B, and C. Clades A and B are further divided into subclades A1, A2, B1, B2, and B3 while clade C had not subclades. Moreover, a clade of strains belonging to the 1962 Fermon reference (GenBank accession AY426531) was present (Genotype US/NL). The dataset contains sequences collected from 1962 to 2022 and the origin of the strains (i.e., collection country) is widespread all over the world (Figure 1). The global circulation of EV-D68 has been dominated by strains belonging to B1 (N=256) and B3 (N=528) subclades. This finding is expected as the B1 and B3 subclade were massively sequenced in Europe, America and China during 2014–2016 for B1 and 2014-2022 for B3. The 94% (N=241) of B1 sequences originated in USA while B3 strains accounting for about 44% of viruses sampled in USA (N=233) while the remaining subgroups were dominated by European samples (37% -Italy: 60, France: 60, Sweden: 54, Netherlands: 25) and China (8% N=43). However, clade A (comprising A1 and A2) account for about 15% (N=149) out of the total strains (N=1027) of which 33% from USA (N=49) and 24% from Europe (N=36). Clade C comprised 41 strains belonging mostly from USA (N=39) while Fermon clade (US/NL genotype) was formed by 23 sequences from USA and France. The Italy strains (N=65) were collected in Lombardy region (Pavia: 30, Milano: 35) in 2022 and grouped into the A2 and B3 subclades representing 3% and 11% of each clade respectively. It has to be noted that the Enterovirus Genotyping Tool was not able to assign genotype for 8 strains (1 Italy, 1 China, 6 USA).

**Figure 3.**
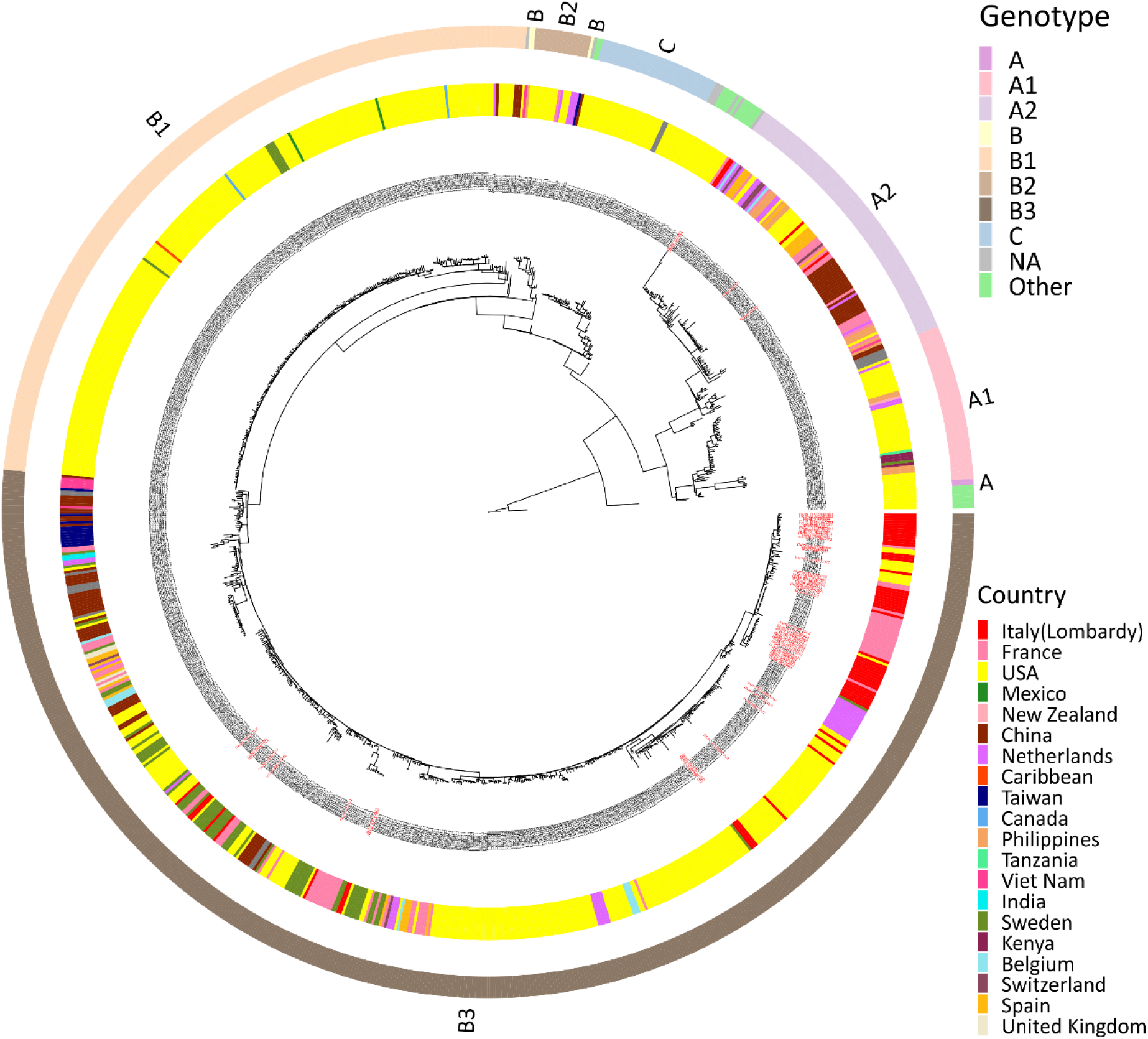
Phylogenetic tree of EV-D68 strains analyzed here and highlighted in red on the tree.

### 3.4. Variability in BC- and DE-loops

VP1 is the major surface protein of the Enterovirus capsid representing the more variable proteins. The protein represents important receptor binding sites and contains hypervariable BC- and DE-loops (i.e., putative neutralizing epitopes). Amino acidic alignment of sequences corresponding to BC- and DE-loop were analyzed for B, B1, B2 and B3 strain (N=806) to calculate the frequency of amino acid changes compared to sequences belonging to Fermon reference (N=3) (Figure 4A). Overall DE-loop remained more conserved with amino acid changes observed in 11 positions out of 14 (78.6%) and in the B3 genotype a great variability was observed. We integrated the phylogenetic tree with the epitope turnover to appreciate the dynamics of the epitope patterns found in the BC- and DE-loops. The antigenic evolution of EV-D68 clearly followed the spread of genotypes. The BC-loop in the A2 subclade, for example, shows 3 prevalent motifs (“KNHTSSEARVDKNFY”, “KNHTSSEARIDKNFY”, “KNHTSSEARTDKNFY”) while A1 clade contains 38 strains out of 53 represented mainly by one motif (“KNHASSEAQTDKNFF”). However, Clade C shares the same prevalent motif of A1 (23 strains) and a new motif previously found in A clade (“KNHASSEARTDKNFF”). Clade B2 is mainly covered by one motif (“KDHTSSAAQTDKNFF”) that is acquired by 23 B3 strains and only one B1 strain. B1 clade shows an abundance of “KDHTSSAAQADKNFF” motif (249 strains) that is prevalent also in B3 clade (219 strains). In the predominantly B3 subgroup a total of 237 out of 528 samples have a BC-loop sequence “KDHTSSTAQTDKNFF” (Figure 4B, dark green point) that is not shared by any of the other clades. The DE loop shows a similar pattern of differentiation as BC-loop with all major clades differing in their sequence (Figure 4B). Clade B1 shows a prevalence of “NGSSNNTYVGLPDL” motifs in 233 strains out of 256. The predominant epitope pattern (“NGSNNNTYVGLPDL”) in the mostly B3 subgroup was present in 254 sequences out of 528 and appear in only one sequence of B1 clade. However, B1 clade prevalent motif (“NGSSNNTYVGLPDL”) is also abundantly present in B3 clade (191/528). The motif evolution patterns revealed in these analyses are agree with the concept of rapid antigenic evolution and epitopes turnover in different genotype strains.

**Figure 4.**
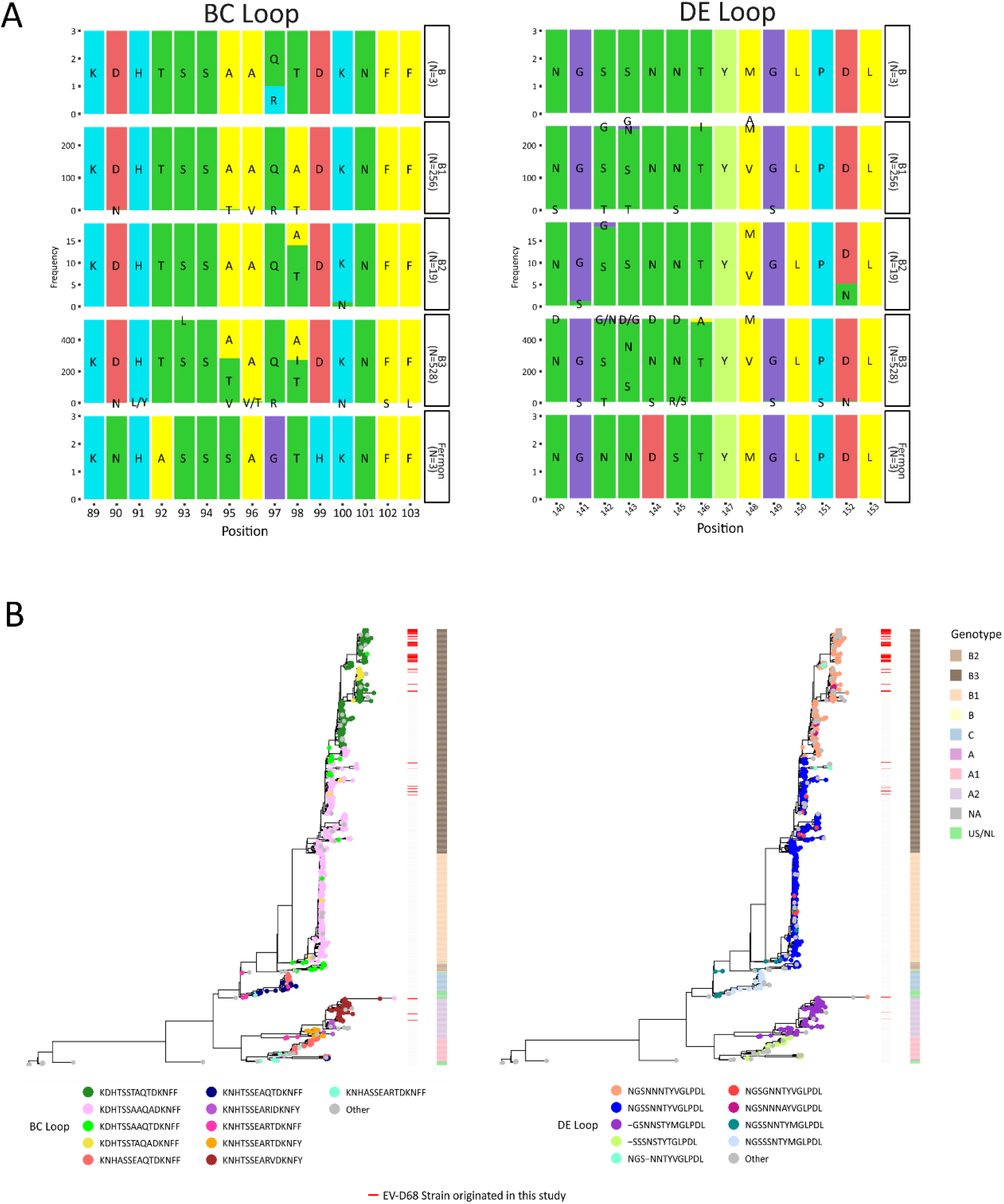
Analysis of the hypervariable BC- and DE-loops by showing amino acid composition (A) or distribution on the phylogenetic tree (B) (BC-loop is amino acid positions 89–103; DE-loop is amino acid positions 140–153).

## 4. Discussion

EV-D68 is an important pathogen in the setting of respiratory viral infections; moreover, it’s able to spread over the nervous system leading to severe neurological syndromes. Thus, molecular surveillance of circulating strains, generally based on VP1gene sequencing, is very needed among high-risk populations, such as children or immunocompromised subjects.

In this study, we designed a new amplicon-based assay for complete genome sequencing of EV-D68 genome. This approach, already used for EV-D68 complete genome sequencing even if with a different number of PCR reactions (Chen *et al*., 2016; Dyrdak *et al*., 2019), consists in a triple PCR reaction to separately amplify three segments of viral genome before molecular libraries preparation and sequencing on Illumina platform. Our PCRs reactions showed to have different performances, with PCR A producing a higher number of amplicons compared to PCR B and C. Overall, sequencing results showed good performances of this newly-design assay. In 73% of all strains with Ct >30 included in the study complete genome sequencing was obtained, with a very good depth of coverage higher than 6000X. Thus, this assay allows a deeper analysis of EV-D68 genetics and molecular evolution not only in the VP1 gene but along the entire genome. On the contrary, in 5% of samples tested, only a partial sequence was obtained, however, the VP1 region was available for further analysis (data not showed). These finding showed that, even if a complete genome sequence was not achieved, VP1 analysis could be performed anyway. Bioinformatic data about EV-D68 complete sequences were analyzed on INSaFLU, a bioinformatic web-based tool that deals with primary sequencing data. Several web-based platforms have been implemented to simply genome reconstruction with a minimum of bioinformatic knowledge. Along with the technical data reported in this study, a phylogenetic analysis on EV-D68 strains collected in 2022 was also performed. All strains belonged to Clade B and 30 amino acid changes were observed in BC-loop and DE-loop regions: these regions are those more exposed on the surface of the viral capsid and the more susceptible to the selective pressure of the immune response.

## 5. Conclusion

The main aim of this study was to design and set up a protocol for whole genome sequencing of EV-D68. Based on the obtained data, this method is robust and reproducible resulting in an easy-to-perform protocol helpful for tracing the evolution of EV-D68 in protein different from VP1. The main commercial kits for whole genome sequencing of viruses are mainly dedicated to the most frequent and studied ones, such as Influenzavirus or Respiratory Syncytial Virus. However, EV-D68 could become a significant pathogen for public health in the next future, and thus this protocol for whole genome sequencing could help clinical and molecular virologists to be ready for molecular epidemiology surveillance.

## Acknowledgments

We thank Daniela Sartori for manuscript editing.

